# L-arabinose induces the formation of viable non-proliferating *Vibrio cholerae* spheroplasts

**DOI:** 10.1101/2020.07.15.203711

**Authors:** Elena Espinosa, Sandra Daniel, Sara B. Hernández, Felipe Cava, François-Xavier Barre, Elisa Galli

## Abstract

A general survival strategy of many life forms faced with harmful growth conditions is to enter into a non-proliferating state until conditions suitable for growth are restored. In bacteria, this survival strategy is associated with antimicrobial tolerance, chronic infections and environmental dispersion. In particular, the agent of the deadly human disease cholera, *Vibrio cholerae*, undergoes a morphological transition from a rod-shaped proliferative form to a spherical non-proliferating form after exposure to cold or cell wall targeting antibiotics. Growth is resumed when the adverse conditions have ceased.

Here, we show that a component of the hemicellulose and pectin of terrestrial plants, L-arabinose, triggers the formation of non-proliferating *V. cholerae* spherical cells, which are able to return to growth when L-arabinose is removed from the growth medium. We found that the cell wall of L-arabinose treated *V. cholerae* cells has a peptidoglycan composition similar to the cell wall of spheroplasts and that they revert to a wild-type morphology through the formation of branched cells like L-forms. Unlike L-forms, however, they are osmo-resistant. Through a random Tn genetic screen for mutants insensitive to L-arabinose, we identified genes involved in the uptake and catabolism of galactose and in glycolysis. We hypothesize that L-arabinose is enzymatically processed by the galactose catabolism and glycolysis pathways until it is transformed in a product that cannot be further recognized by *V. cholerae* enzymes. Accumulation of this enzymatic by-product triggers the formation of viable non-dividing cell wall deficient spherical cells.

## Introduction

Cholera is an acute diarrhoeal disease caused by ingestion of food or water contaminated with *Vibrio cholerae*, a curved rod shape bacterium that propagates in warm briny and salty waters. Cold stops *V. cholerae* proliferation. However, the bacterium has the ability to persist for months in cold water under the form of a spheroplast, i.e. a cell wall deficient spherical cell, and return to growth when the sea temperature rises [1–3]. Similarly, *V. cholerae* is known to persist under a spherical form in biofilms [4]. Correspondingly, it was observed that *V. cholerae* tolerates exposure to antibiotics inhibiting cell wall synthesis [5].

The high propagation rate of *V. cholerae* and its capacity to survive unfavourable growth conditions have led to several pandemics, which have caused and are still causing major socio-economic perturbations [6]. Seven cholera pandemics have been recorded since the beginning of the 18^th^ century. The 7^th^ pandemic started over 50 years ago and is still ongoing. *V. cholerae* strains can be grouped into 12 distinct lineages, out of which only one gave rise to pandemic clones [7]. Transition from the previous (6^th^) to the current (7^th^) pandemic was associated with a shift between 2 different phyletic subclades, the so-called classical and El Tor biotypes [7–11]. Isolates of the current pandemic are rapidly drifting and it is suspected that the constant appearance of new atypical pathogenic variants of *V. cholerae* will eventually lead to a more virulent strain that will start a new (8^th^) pandemic. This motivated extensive research on the physiology of the bacterium and its evolution towards pathogenicity [7–11].

Inducible promoters permit to study the phenotypes of null mutations of essential genes and/or to trigger the production of potentially toxic recombinant protein fusions at specific stages of the cell cycle. In particular, a tightly-regulated high-level expression inducible system based on the promoter of the *Escherichia coli araBAD* operon and its regulator *araC*, known as the P_BAD_ system [12], was and still is instrumental to many *V. cholerae* studies. The AraBAD enzymes allow *E. coli* to exploit L-arabinose (L-Ara), a component of the hemicellulose and pectin of terrestrial plants, as a carbon and energy source [13]. AraC acts both as a positive and a negative regulator, repressing P_BAD_ in the absence of L-Ara and activating its transcription when bound to it [13]. *V. cholerae* lacks a *bona fide* arabinose import and metabolization pathway. Nevertheless, the *E. coli* P_BAD_ system proved to be very effective in *V. cholerae*, which suggested that L-Ara was imported in the cytoplasm of the cells. However, we and others recently reported that L-Ara could interfere with the growth of the bacterium, calling for a better understanding of the impact of L-Ara on the physiology of *V. cholerae* and the pathway leading to its toxicity [14].

Here, we show that *V. cholerae* cells stop dividing or elongating and lose their characteristic curved rod cell shape in the presence of >0.5% (w/v) and >0.1% (w/v) of L-Ara in rich and poor media, respectively. *V. cholerae* cells become spherical and morphologically akin to the spheroplasts obtained by exposure to cold temperatures [1,2,15] or cell wall targeting antibiotics [5]. We found that the spheroplasts are able to revert to exponentially growing rods in only a few generations when L-Ara is removed, demonstrating that they remain viable. We further show that L-Ara is imported by the galactose transport system and that it interferes with the metabolism of *V. cholerae* by entering the galactose and glycolysis catabolic pathways. Finally, we show that *V. cholerae* mutants with impaired physiology are more sensitive to the presence of L-Ara, morphologically transitioning to spheroplasts with as little as 0.01% (w/v) of L-Ara in poor media. Taken together, these results suggest that spheroplasts formation might be a general survival strategy of *Vibrios* when faced with conditions that perturb their metabolism. From a technical point of view, they indicate how the P_BAD_ expression system can still be used in *V. cholerae* without perturbing the metabolism of the bacterium.

## Results

### L-arabinose induces the formation of non-dividing spherical cells

To study the effect of L-Ara on cell morphology and growth, we added increasing concentrations of L-Ara to wild-type N16961 *V. cholerae* cells under exponential growth in different liquid media. Cells grown in M9-MM appeared with a wild-type rod shape in the absence or up to 0.02% (w/v) L-Ara, but the entire cell population became spherical in the presence of 0.1% (w/v) L-Ara (Figure 1A). In cultures grown in M9-MM supplemented with casamino acids (CAA), spherical cells started to appear at 0.2% (w/v) of L-Ara. At 0.5% (w/v) of L-Ara, all cells exhibited a spherical morphology. In LB, the concentration of L-Ara had to be increased to 1% (w/v) to induce loss of the normal cell shape (Figure1A). The morphological appearance of L-Ara treated cells resembles that of non-proliferative cells obtained by incubating *V. cholerae* cells at 4°C [2,4] or by treating them with cell wall targeting antibiotics [5] (Figure 1B).

**Figure 1.**
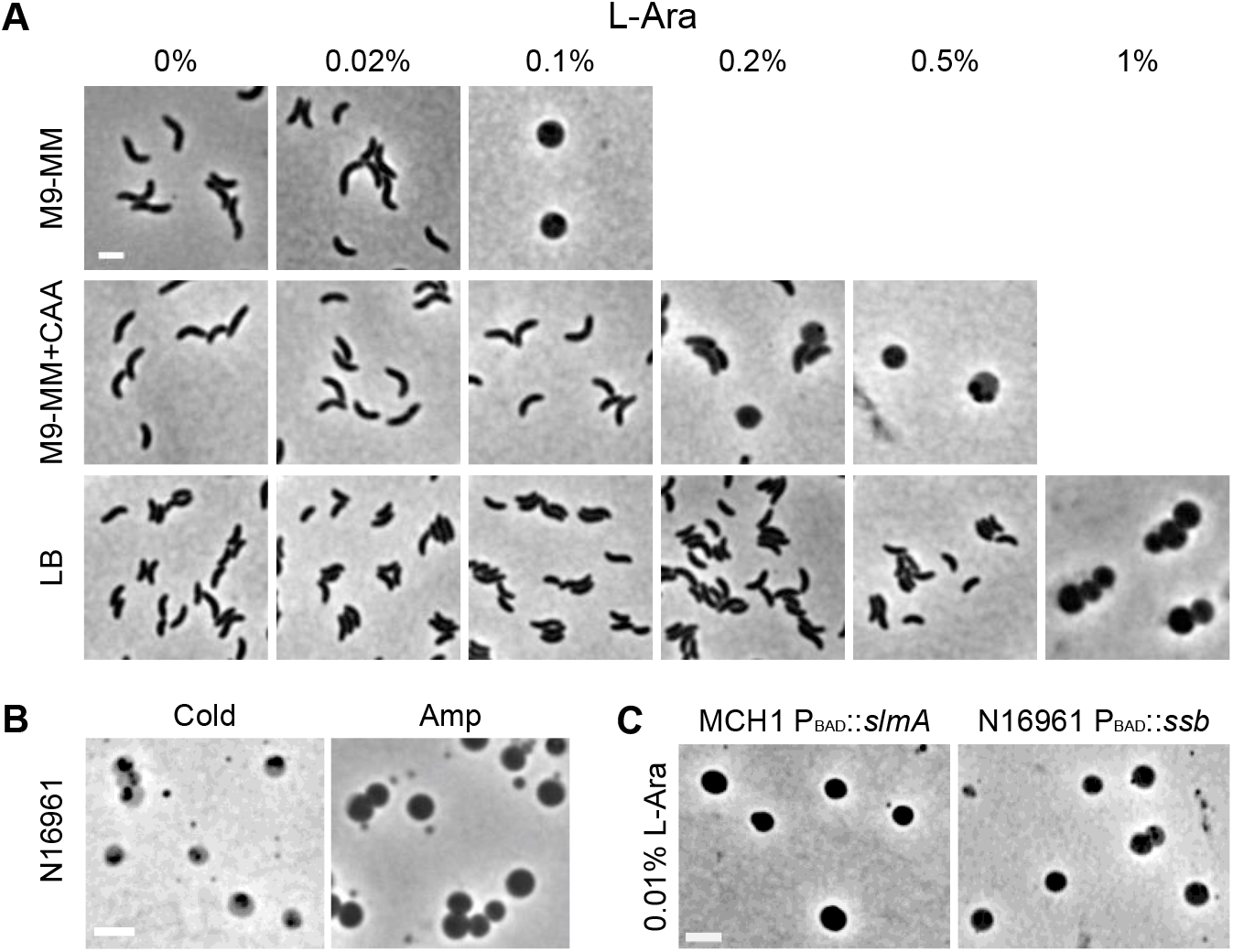
L-Ara induces loss of rod shape. **A.** Phase contrast images of *V. cholerae* N16961 cells grown at 30°C in the indicated media with increasing concentrations of L-Ara. **B.** Phase contrast images of *V. cholerae* N16961 cells incubated in M9-MM at 4°C for 6 weeks (left panel) and grown in LB with 100 μM Ampicillin (Amp) at 30°C (right panel). **C.** Phase contrast images of *V. cholerae* EGV299 (MCH1 P_BAD_*::YGFP-slmA*) and EGV300 (N16961 P_BAD_::ssb-YGFP) cells grown in M9-MM at 30°C in presence of 0.01% (w/v) L-Ara. Scale bars = 2 μm.

In parallel to microscopic inspection, we followed the optical density of cell cultures over time. In all tested media, L-Ara had a detrimental effect on cell proliferation at the same concentrations at which loss of cell morphology was observed (Supplementary Figure 1). Additional sugars or carbon sources were tested for similar effects on cell shape and growth, but none of them, including D-arabinose, affected *V. cholerae* growth or morphology (Table 1 and Supplementary Figure 2).

**Table 1.** Carbon sources tested for N16961 rod shape loss. They were added to M9-MM at a concentration of 0.2% (w/v), with the exception of glycerol at 10 % (v/v).

| Carbon source | Growth | Spherical cells |
| --- | --- | --- |
| D-Arabinose | + | — |
| L-Arabinose | — | + |
| L-Rhamnose | + | — |
| Glucose | + | — |
| Galactose | + | — |
| Glycerol | + | — |
| Sucrose | + | — |

### Mutants with impaired physiology are more sensitive to L-Ara

Strikingly, we observed that physiologically impaired *V. cholerae* mutants were more sensitive to the presence of L-Ara. In particular, the presence of as little as 0.01% (w/v) of L-Ara was sufficient to induce the formation of non-dividing spherical cells in a N16961 strain carrying two copies of the *ssb* gene, which codes for an essential single strand DNA binding protein implicated in the regulation of replication, transcription and homologous recombination repair [16] (Figure 1C). Likewise, the presence of 0.01% (w/v) of L-Ara was sufficient to induce the formation of non-dividing spherical cells in MCH1, a mono-chromosomal *V. cholerae* strain, when the SlmA nucleoid occlusion protein was overproduced [17] (Figure 1C).

### Transition dynamics to spherical cells at the population level

To visually inspect the morphological transition at the population level over time, we collected cell samples every hour for 10 hours after L-Ara addition and examined them at the microscope. Cells were divided in three categories based on their shape: cells with a rod shape, cells composed of a rod and a small or large irregular bulge protruding from the cell wall, which we refer to as bleb, and cells with a spherical shape (Figure 2A). Spherical cells started to appear after 5 hours and comprised 90% of the cell population after 9 hours. The sharp increase in the proportion of spherical cells in the population corresponded to an equally fast decline in rod shaped cells, which dropped to less than 10% of the cell population at the end of the experiment. Cells composed of a rod and a bleb appeared around 4 hours after L-Ara addition. Blebs were randomly located on the surface of the cells. In particular, there was no preference for mid-cell or cell pole locations (Supplementary Figure 3). Cells with protruding blebs never represented more than 5% of the entire cell population and almost disappeared at the end of the time course, which suggested that they corresponded to a transient state between the rod and the spherical state (transitioning cells). Taken together, these results suggest that a few hours are required after the growth arrest before morphological transition. However, once transition is initiated the formation of spherical cells is very fast.

**Figure 2.**
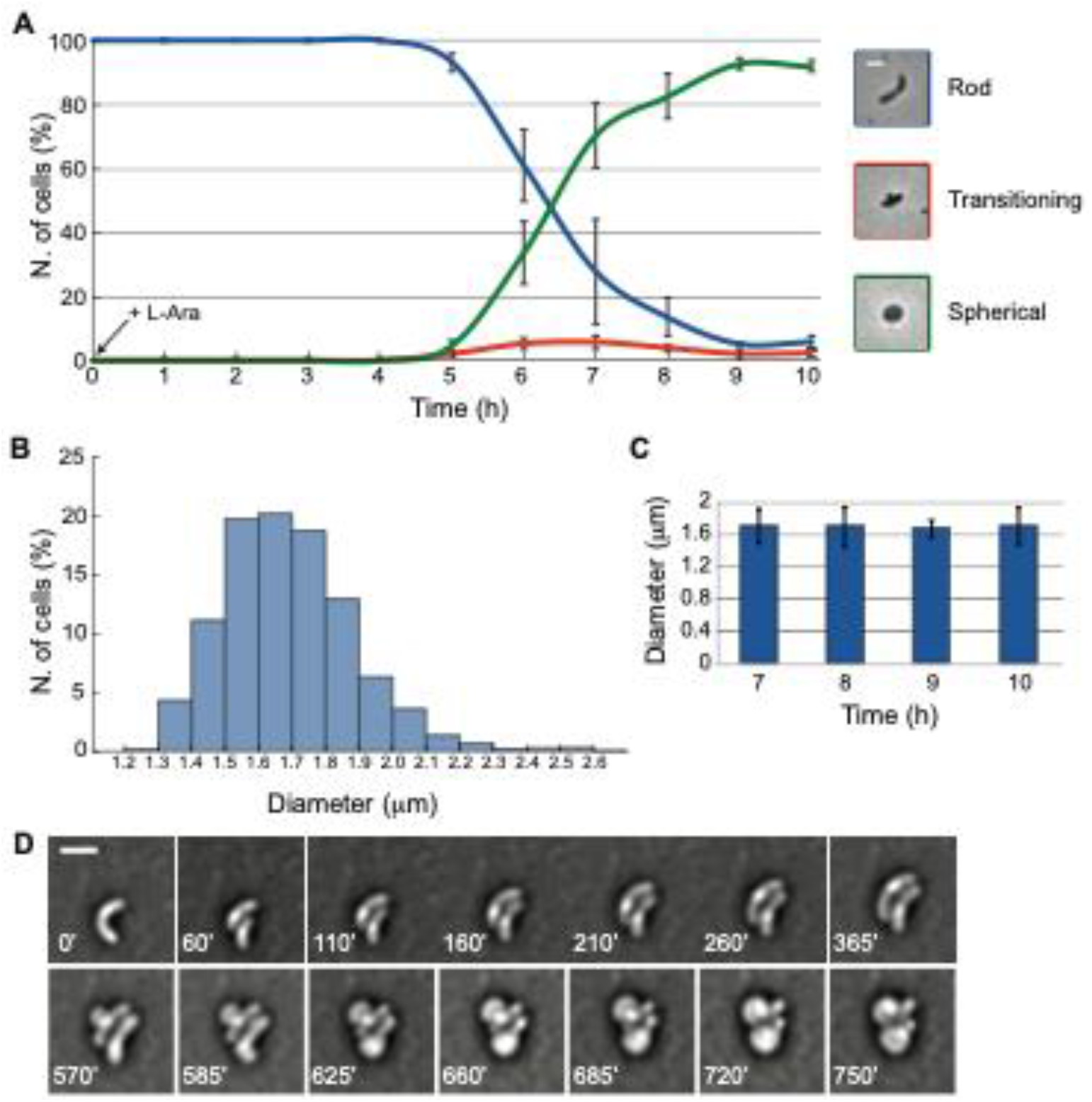
Transition dynamics to spherical cells. Cells were grown in M9-MM at 30°C. L-Ara was added at a concentration of 0.2% (w/v) when indicated. **A.** Kinetics of *V. cholerae* morphological change from rods to spherical cells. Cell shape was inspected at the microscope every hour after L-Ara addition. A representative image for each cell category (rod, transitioning, spherical) is represented. Mean of three independent replicates and the standard deviation are represented. **B.** Diameter distribution of *V. cholerae* spherical cells treated with L-Ara for 10 hours. **C.** Average diameter of *V. cholerae* spherical cells over time. Mean of three independent replicates and the standard deviation are represented. **D.** Transition from rods to spherical forms. N16961 cells were mounted on a M9-MM agarose pad containing 0.2% (w/v) L-Ara. Bright-field still images from time-lapse microscopy experiments. Images were taken every 5 minutes for 12.5 hours. Scale bars = 2 μm.

The size of the spherical cells was heterogeneous (Figure 1). The diameter of the majority of cells was comprised between 1.5 and 1.8 μm (Figure 2B). The average of the diameter did not change over time, suggesting that spherical cells neither decreased nor increased in volume after their formation (Figure 2C). Based on the average cell diameter of spherical cells and the average cell length and width of rod-shaped cells, we estimate that the cell volume of spherical cells is around 2.5 times bigger than that of exponentially growing rod cells.

### Transition dynamics to spherical cells at the single cell level

We performed time-lapse video-microscopy experiments to inspect the transition process from rod to sphere at the single cell level (Figure 2D and Movie 1). All the observed cells displayed the same transition pattern. After exposure to L-Ara, a single bleb appeared at the bacterial cell surface. As the bleb increased in size, the original cell was assimilated into the forming sphere until the original rod shape was completely lost. The time when a bleb became visible on the cell surface varied from cell to cell. However, once started, completion of the morphological change was comparable in all the cells. On an agarose M9-MM pad, blebbing cells transitioned to spheres in around 2 to 3 hours. We could observe rare events in which cells failing to transition to spheres lysed (Movie 2).

### L-Ara induced spherical cells are cell wall deficient

The spherical shape of L-Ara treated cells reminded that described for cell wall deficient (CWD) forms that have completely (protoplast-type) or almost entirely (spheroplast-type) lost the peptidoglycan (PG) layer, which suggested a process of cell wall degradation or PG remodelling mechanism [18]. To verify this point, we compared the PG content and composition of exponentially growing *V. cholerae* cells and L-Ara induced non-dividing spherical cells (Table 2). To limit contamination by the PG of cells that had not completely transformed into spheres in the presence of L-Ara, we used MCH1 cells expressing an additional copy of SlmA because of their increased sensitivity to L-Ara (Figure 1C).

**Table 2.**
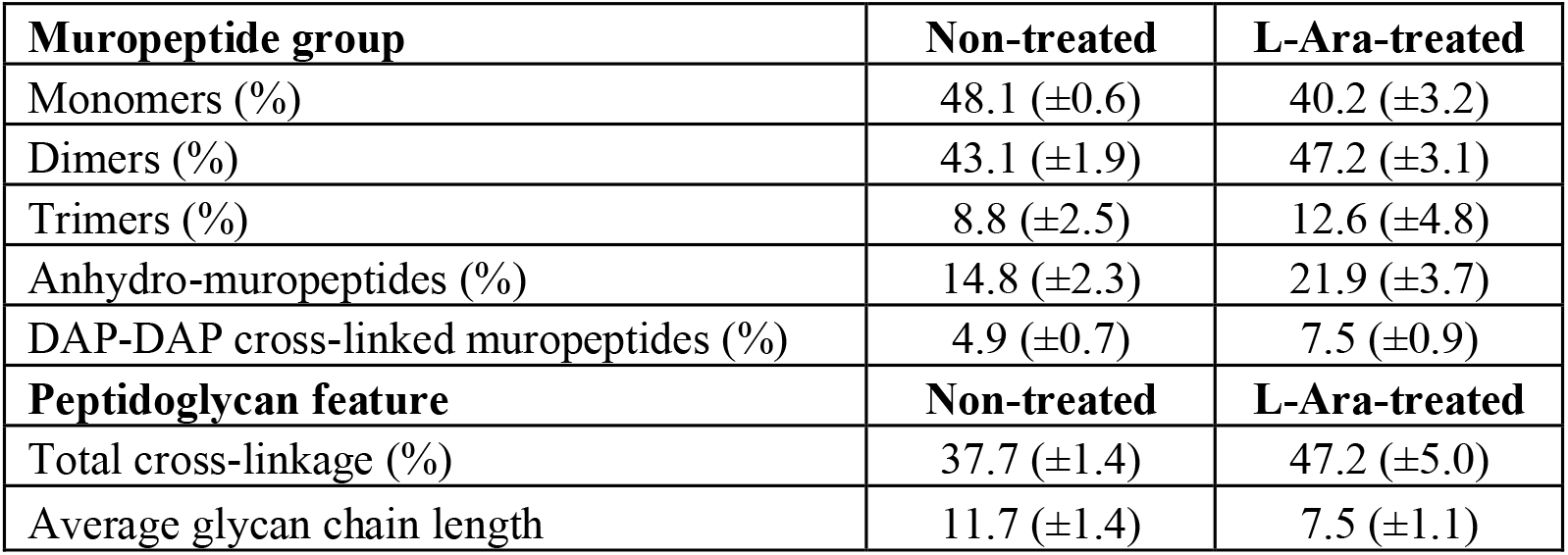
Quantification of muropeptides, peptidoglycan cross-linking levels and average chain length of L-Ara treated and non-treated cells. Values are the means of three independent experiments and the standard deviation is represented.

The PG of exponentially growing and L-Ara treated cells that had completely transitioned to spheres was extracted and submitted to UPLC analysis (Supplementary Figure 4A). The area of the UPLC profiles showed that the amount of PG per cell in L-Ara treated bacteria was around 10 times lower than the amount present in rod-shaped cells (Supplementary Figure 4B). In addition, we found that glycan chains were twice shorter in L-Ara treated cells than in untreated cells. The length of glycan chains is calculated based on the number of anhydro-muropeptides [19], which result from the activity of lytic transglycosylases [20,21]. Therefore, a reduction in the average glycan chain length suggests an increase of lytic transglycosylase activity in the presence of L-Ara. Finally, we observed that the amount of DAP-DAP cross-linked muropeptides significantly increased in L-Ara treated cells. Taken together, these results indicate that L-Ara induced spherical cells are spheroplasts.

### L-Ara induced spheroplasts are viable and can revert to proliferation

To evaluate if L-Ara had a detrimental effect on cell viability we estimated the number of viable bacteria in a time course experiment. An initial culture was split in two after 2 hours of growth and L-Ara added to one of the two halves. The number of viable cells in the cultures was determined by plating aliquots on LB plates and counting the numbers of colonies. Viable cell count kinetics (represented as colony forming units, CFU) showed that L-Ara addition had an immediate inhibitory effect on cell proliferation but did not cause a corresponding decline in cell viability (Figure 3A). Indeed, 75% of the number of cells before L-Ara addition gave rise to colonies after an incubation of 10 hours with L-Ara (Figure 3B).

**Figure 3.**
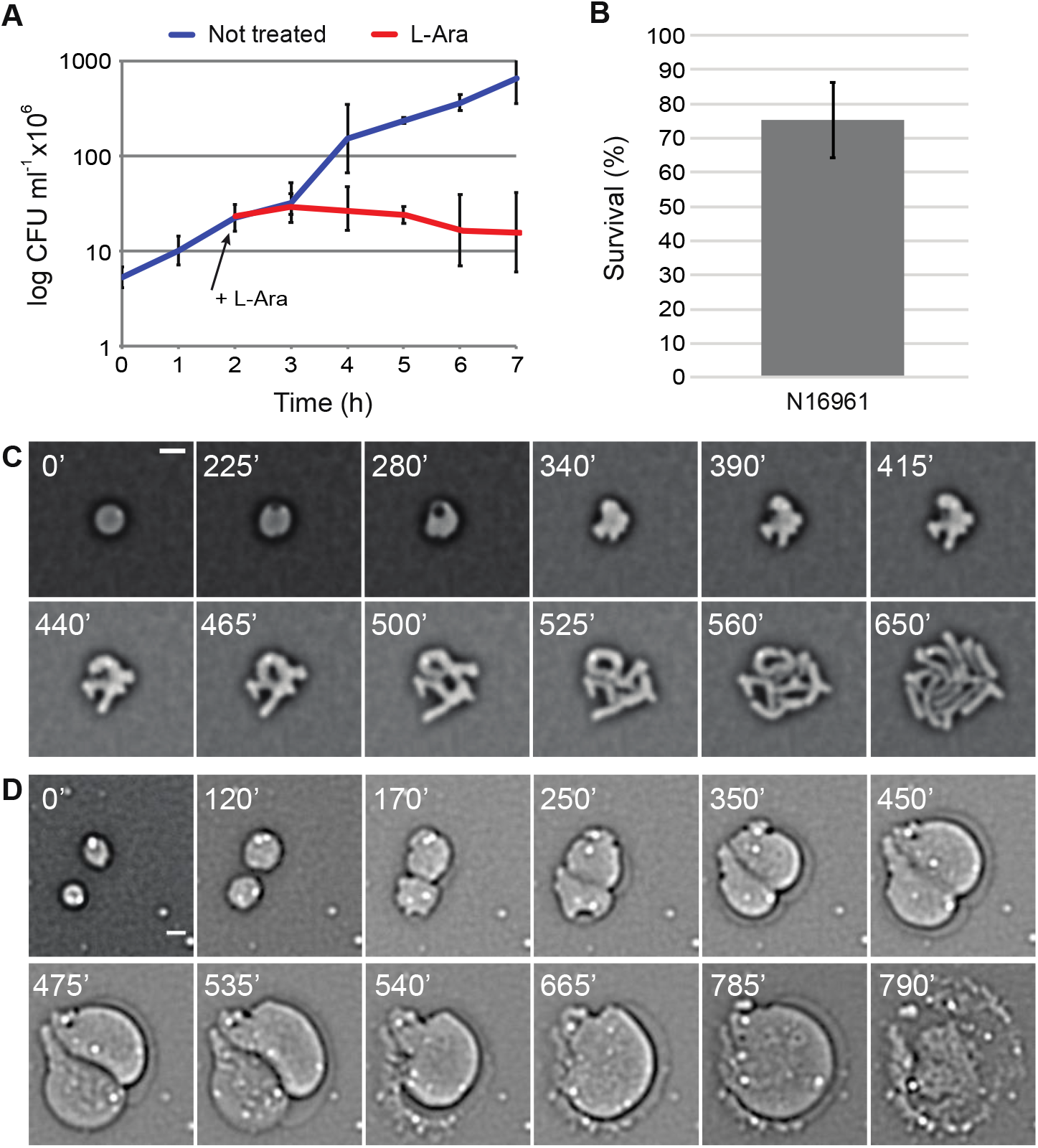
Recovery of growth and rod shape. Cells were grown in M9-MM at 30°C. L-Ara was added at a concentration of 0.2% (w/v) when indicated. Viable colony count (CFU) of *V. cholerae* cells grown with and without L-Ara over time (**A**) and after 10 hours (**B**). Mean of three independent replicates and the standard deviation are represented. In the time-lapse experiments, cells were grown in M9-MM + 0.2% (w/v) L-Ara until they became spherical and then mounted on a M9-MM agarose pad in the absence of L-Ara. Bright-field still images were taken every 5 minutes for 14 hours. **C**. N16961 cells recovery from spherical CWD forms to rods. **D.** Spherical *wigKR* cells are not able to recover rod shape. Scale bars = 2 μm.

Time-lapse video-microscopy was performed to determine how non-proliferating spherical cells could return to proliferation and recover a curved rod shape after L-Ara removal. Single cell analyses showed that reversion to proliferating rods started with the elongation of the spheroplasts, which was followed by the formation of multiple protrusions on their surface (Figure 3C and Movie 3). The protrusions elongated outward, giving rise to multiple branched cells, and the original rod shape was recovered after only a few division events. The time required to initiate elongation greatly differed from cell to cell: the recovery process of some cells started almost immediately after the removal of L-Ara and took a few hours for other cells. However, the length of time between the initiation of the recovery process and its completion was similar for all the cells. On an agarose M9-MM pad, cells transitioned from elongating spheres to symmetrically dividing rods in around 4 to 5 hours.

Time-lapse video-microscopy observations further suggested that the overall 25% loss of CFU after 10 hours of L-Ara treatment (Figure 3B) was accounted for by the number of cells that lysed during the transition to spheroplasts after the addition of L-Ara (Movie 2) and by those that lysed during the recovery process after L-Ara removal (Movie 4). As no osmo-protectant was added to the growth media, this observation suggests that the spheroplasts are osmo-resistant.

### *wigKR* is essential for the recovery of cells after L-Ara treatment

The histidine kinase/response regulator pair *wigKR* is thought to induce a higher expression of the full set of cell wall synthetic genes in response to cell wall damages [22]. It was previously reported that *wigKR* was essential for the recovery of the cell shape and the return to proliferation of *V. cholerae* cells treated with cell wall targeting antibiotics [22]. In the presence of L-Ara, *wigKR* cells formed spheroplasts at the same timing than wild-type *V. cholerae* cells and blebs remained distributed all around the surface of cells (Supplementary Figure 5A). However, reversion was dramatically different. After removal of L-Ara, *wigKR* spheroplasts immediately started to grow in diameter, expanding continuously in size until they exploded (Figure 3D and Movie 5). On the contrary, *wigKR* cells that had stopped dividing but had not yet transitioned to spheroplasts were able to return to a proliferative state without any obvious defect (Movie 6). After 10 hours incubation with L-Ara, only 10% of the *wigKR* cells could still form colonies (Supplementary Figure 5B). It corresponded to the proportion of *wigKR* cells that did not transition to spheres upon L-Ara addition, suggesting that *wigKR* was essential for the recovery of the spheroplasts, as previously observed for cells treated with cell wall targeting antibiotics.

### L-Ara is imported in the cytoplasm and metabolically processed

We reasoned that metabolic arrest and spheroplast formation were unlikely to result from the action of L-Ara on the surface of *V. cholerae* or in its periplasm, in contrast to cell wall targeting antibiotics. *V. cholerae* lacks a *bona fide* arabinose import and metabolization pathway. However, the effectiveness of the *E. coli* P_BAD_ promoter regulation by L-Ara suggested that it was at least passively imported in the cytoplasm of *V. cholerae* cells. To identify putative factors involved in the response of *V. cholerae* cells to L-Ara, we screened a random transposition (Tn) library for mutants that were insensitive to L-Ara. To limit false positives, the screening was performed using the MCH1 strain carrying an additional copy of the *slmA* gene that fully transitions to spheres in the presence of as little as 0.01% (w/v) of L-Ara (Figure 1C). After Tn mutagenesis, the libraries were screened on M9-MM plates containing 0.1% (w/v) L-Ara. We then isolated single L-Ara resistant colonies on fresh L-Ara plates and verified their resistance to L-Ara in liquid media. Finally, the morphology of the mutants treated with L-Ara was inspected at the microscope.

Of the 11 colonies initially detected in the screening, 9 were confirmed to be L-Ara insensitive. They corresponded to Tn insertions in 6 different genes (Table 3 and Supplementary Figure 6). The suppressor capacity of the genes identified in the screening was confirmed by using the corresponding mutant in an ordered *V. cholerae* mapped Tn library [23] or, if the strain corresponding to the inactivated gene of interest was missing (i.e. *vc0263*, *vc2689*), by direct mutagenesis of the gene in a N16961 wild-type background. Comparison of the growth rates of the identified mutants in the presence or absence of L-Ara further confirmed that they were not affected by L-Ara (Supplementary Figure 7). The study was completed by the investigation of the resistance to L-Ara of a few genes belonging to the same enzymatic pathways of the mutants identified in the screening.

**Table 3.** Suppressor mutants of L-Ara induced toxicity identified in a Tn-based screen.

| **Gene** | **Putative function** | **N. hits** |
| --- | --- | --- |
| *vc0263* | Galactosyl-transferase | 2 |
| *vc1325* | Galactoside ABC transporter, periplasmic D-galactose/D-glucose binding protein | 1 |
| *vc1327* | Galactoside ABC transporter, ATP-binding protein | 1 |
| *vc1328* | Galactoside ABC transporter, permease protein | 1 |
| *vc1595* | Galactokinase | 1 |
| *vc2689* | 6-Phosphofructokinase, isozyme I | 3 |

Inactivation of *vc1325*, *vc1327*, *vc1328*, *vc1595*, *vc1596* or *vc0263* fully restored growth in presence of L-Ara (Figure 4). VC1325, VC1327 and VC1328 are homologues of *E. coli* MglB, MglA and MglC, respectively. The three corresponding genes form an operon encoding the components of the ABC galactose transport system: MglB is the periplasmic binding protein, MglA the ATP-binding protein and the integral membrane protein MglC is the permease [24,25]. VC1595 is the homologue of the *E. coli* GalK galactokinase, the first enzyme in the Leloir pathway of galactose metabolism [26]. VC1596 is the homologue of *E. coli* GalT, the galactose 1-phosphate uridylyltransferase. VC0263 is a homologue of *E. coli* WcaJ, the initiating enzyme for colanic acid synthesis [27], which was also described to act as a galactose-1-phosphate transferase *in vitro* [28]. Taken together, these results suggest that L-Ara is imported in the cytoplasm of *V. cholerae* by the galactose transporter and processed by the galactose catabolic enzymes. In *Sinorhizobium meliloti*, the arabinose transporter AraABC has been described to play a role in galactose uptake [29], suggesting a similarity in the activity of the arabinose and galactose transporter.

**Figure 4.**
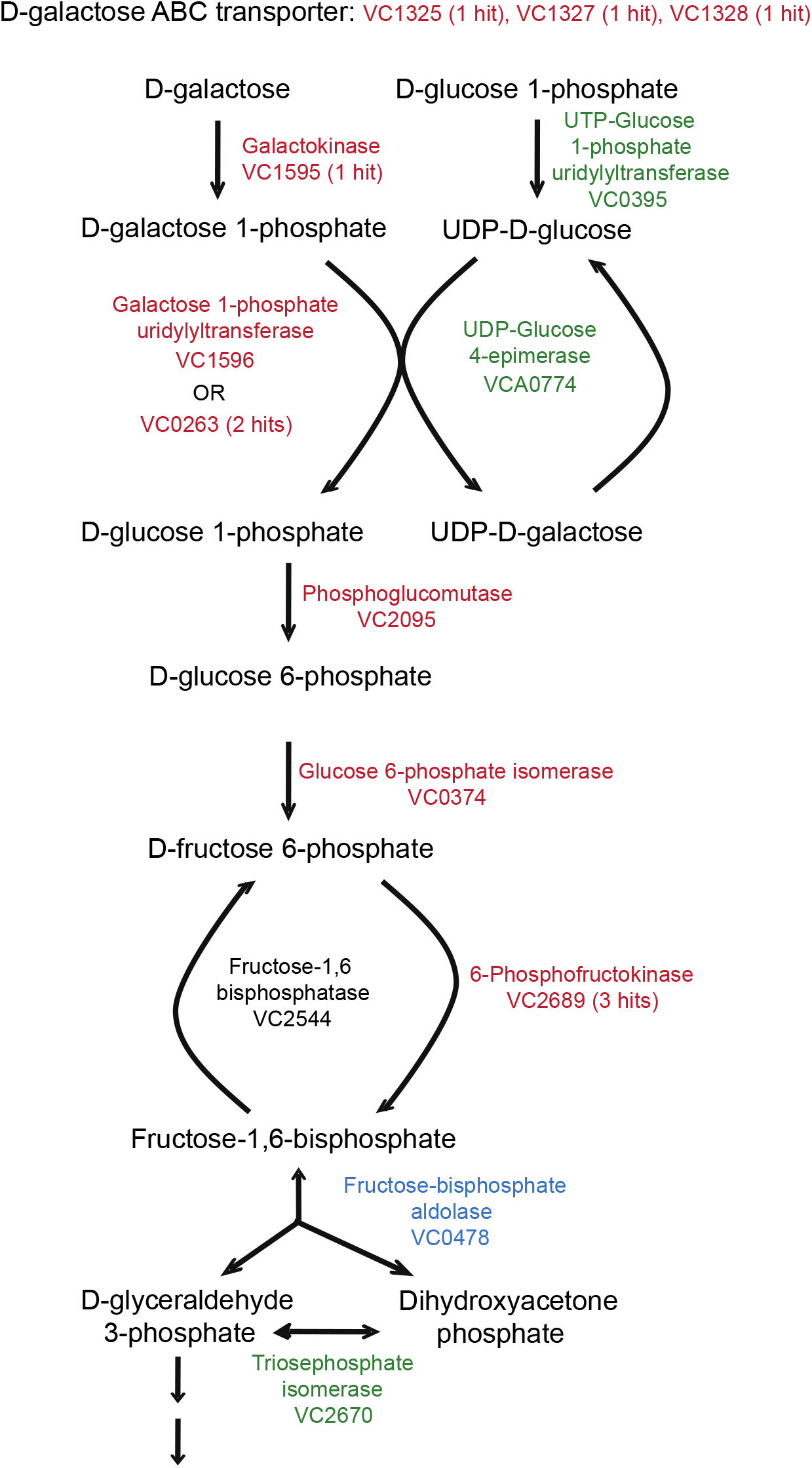
L-Ara insensitive mutants. Schematic representation of the enzymatic pathways targeted by the suppressor screening. In red are the genes that when inactivated suppress the L-Ara induced phenotype (number of hits obtained in the screening in between parentheses), in green the non-suppressors, in blue the gene that could not be inactivated because essential and in black the gene not tested.

Growth of a *vc2095* mutant was very slow, and further decreased in the presence of L-Ara. However, its growth was not completely arrested and spherical cells were not detected. VC2095 is the homologue of *E. coli* Pgm phosphoglucomutase, which suggested that the product resulting from the action of the galactose catabolic enzymes on L-Ara entered the glycolysis pathway. Mutants of *vc0374*, the homologue of the *E. coli* glucose 6-phosphate isomerase *pgi*, grew very slowly and L-Ara addition did not further worsen the growth defect. Microscopic inspection of L-Ara treated cells showed the presence of rare spherical cells. In addition, three independent Tn hits were obtained in VC2689, the homologue of *E. coli* 6-phosphofructokinase PfkA. L-Ara reduced the growth rate of *vc2689* mutants (Supplementary Figure 7). However, microscopic inspection revealed a majority of rod-shaped cells with a few isolated spherical cells, suggesting that L-Ara sensitivity was reduced even though not completely suppressed.

The genes involved in the successive reactions of glycolysis were essential and could not be inactivated, with the exception of *vc2670*, the putative homologue of *E. coli* triosephosphate isomerase *tpiA*, which did not suppress L-Ara induced growth inhibition (Supplementary Figure 7). Finally, we found that Tn inactivation of *vc0395* and *vca0774*, the homologues of *E. coli* UTP-glucose 1-phosphate uridylyltransferase GalU and UDP-glucose 4-epimerase GalE, respectively, which are necessary for supplying UDP-glucose and the interconversion of UDP-galactose as part of the galactose metabolism [30], did not suppress L-Ara induced growth inhibition (Supplementary Figure 7).

Taken together, these results suggest the growth arrest of *V. cholerae* cells treated with L-Ara and their transition to spheroplasts is due to the entrance of L-Ara in the galactose and glycolysis pathways.

## Discussion

It was recently reported that L-Ara inhibits the proliferation of *V. cholerae* [14]. Here, we show that it is associated with a change in the morphology of the cell from a curved rod shape to a spherical form (Figure 1). The spherical cells lose the ability to divide and stop growing in volume over time, suggesting a major metabolic arrest (Figure 2). We found that L-Ara induced spherical cells are spheroplasts, i.e. they are cell wall deficient (Table 2 and Supplementary Figure 4). Nevertheless, they remain viable and once L-Ara is removed from the environment they resume proliferation and revert to the original cell shape after a few divisions (Figure 3).

### Metabolism arrest is linked to the processing of L-Ara by the galactose pathway

*V. cholerae* is known to be sensitive to high concentrations of several carbon sources, including glucose, in starvation conditions [31]. In contrast, L-Ara can stop *V. cholerae* proliferation in both fast and slow growth conditions (Figure 1). The phenomenon was strictly limited to L-Ara (Table 1). *V. cholerae* lacks a *bona fide* L-Ara import and degradation pathway. However, the effectiveness of the regulation of the *E. coli* P_BAD_ promoter in *V. cholerae* indicated that it was at least passively imported in the cytoplasm of the bacterium, where it interfered with the metabolism. We performed a Tn-based screening to determine which cellular processes might be involved in the action of L-Ara. The majority of the genes identified belonged to the galactose Leloir and glycolysis pathways, starting from the uptake of L-Ara through the ABC galactose transport system (Table 3). The distribution of suppressors all along the galactose catabolic pathway and the first steps of glycolysis strongly supports the hypothesis that L-Ara is mistakenly recognized as a substrate of the galactose Leloir pathway and that through a series of enzymatic reactions it is converted into a product that enters the glycolysis pathway (Figure 4). The last enzyme in the glycolytic pathway we identified in the Tn suppressor screening is the putative homologue of 6-phosphofructokinase. These results suggest that L-Ara is converted into a phosphorylated sugar by-product that cannot be further metabolized. We found that two enzymes can perform the last step of the Leloir pathway on L-Ara, the Galactose 1-phosphate uridylyltransferase (VC1596) and the product of *vc0263*. Inactivation of either *vc1596* or *vc0263* was sufficient to make *V. cholerae* insensitive to L-Ara, further suggesting that growth arrest is not directly triggered by the non-metabolizable phosphorylated sugar by-product of L-Ara but by the metabolic imbalance created by its accumulation. Correspondingly, L-Ara arrests growth all the more efficiently in slow growing conditions, i.e. in conditions where the concentration of L-Ara is high compared to other nutrients or in cells carrying mutations that already affect their physiological state (Figure 1). Several other studies suggested that accumulation of a phosphate ester metabolite could perturb growth: L-Ara inhibits the growth of *E. coli araD* mutants because of the accumulation of L-ribulose 5-phosphate [32]; Galactose inhibits the growth of *E. coli galT* mutants because of the accumulation of galactose 1-phosphate [33,34]; Rhamnose stops the growth of *Salmonella* Typhi strains defective in the rhamnose degradation pathway because of the accumulation of L-rhamnulose 1-phosphate [32,35].

### L-Ara mediated metabolic perturbation does not prevent the use of P_BAD_

Importantly, the realization that L-Ara can perturb the metabolism of *V. cholerae* does not jeopardize previous results obtained with the *E. coli* P_BAD_ expression system in this bacterium since L-Ara concentrations lower than those that promote growth arrest and spheroplasts formation were almost always used (Figure 1). However, our study indicates that special care should be taken in future works when using the *E. coli* P_BAD_ expression system in mutants of *V. cholerae* (Figure 1). According to our results, two methods can be proposed to avoid metabolic artefacts when using the P_BAD_ expression system in *V. cholerae*. On one hand, cells can be grown in media containing glucose, which represses expression of the galactose import and degradation pathway [34]. On the other hand, experiments can be performed in cells that can import L-Ara but are insensitive to it by mutating the *galK* gene, which codes for the first enzyme that processes L-Ara in the cytoplasm (Figure 4).

### Spheroplasts formation results from an imbalance in the cell wall maintenance enzymes

It was reported that *V. cholerae* cells tolerate exposure to antibiotics inhibiting cell wall synthesis by transitioning to non-dividing cell wall deficient spherical cells [5]. Like L-Ara induced spheroplasts, those spherical cells are osmo-resistant and can revert to proliferating rods after removal of the antibiotics [5]. However, in contrast to L-Ara induced spheroplasts, *V. cholerae* cells exposed to antibiotics inhibiting cell wall synthesis grow in volume over time, suggesting that they are not dormant (Figure 3, [36,37]). In this regard, antibiotic treated cells are more similar to bacterial L-forms, which can be obtained in osmotic stabilizing media in several microorganisms, including *E. coli*, by treating cells with lysozyme [38], by adding the β-lactam cefsulodin (a specific inhibitor of the penicillin binding proteins PBP1A and PBP1B) [39,40], or by inhibiting synthesis of the Lipid II cell wall precursor with fosfomycin [41]. The series of steps and the proteins involved in the formation of antibiotic induced spherical cells in *V. cholerae* have been partially elucidated [5]. The PG amidase AmiB is responsible for the first breach in the cell wall from which blebs originate, although other enzymes can initiate the process in its absence. For the transition from blebs to viable spheres either the D,D-endopeptidase ShyA or ShyC are required, otherwise cells lyse after blebbing. At last, lytic transglycosylases appear to play a role in cell wall degradation and PG turnover and recycling. The analysis of the muropeptide composition of L-Ara treated *V. cholerae* cells showed that they still maintained a residual amount of PG whose structure was remarkably similar to that described for *E. coli* cefsulodin-induced L-forms [39]. The dramatic decrease in the average chain-length and corresponding increase in anhydro-muropeptides hint to a higher activity of lytic transglycosylases in cleaving the PG and producing shorter chains. The increase in DAP-DAP cross-linkage, an unusual kind of cross-linkage specifically generated by L,D-transpeptidases [42], further suggests that the metabolic arrest induced by L-Ara inhibits PBP enzymes and stimulates the L,D-transpeptidase activity (LdtA [43]). Taken together, these results suggest that L-Ara promotes the formation of spheroplasts because it induces a metabolic arrest that leads to an imbalance between PG synthesis and degradation. In this regard, L-Ara treatment mimics the action of cell wall targeting enzymes. However, mid-cell does not appear to be the preferential site for bleb formation in L-Ara treated cells as described for cells exposed to antibiotics targeting cell wall synthesis. Therefore AmiB, which is supposedly active only at the septal site [44,45], might not be the preferred initiator of cell wall cleavage in the presence of L-Ara.

### Proliferation restart results from the restoration of a normal metabolism

*wigKR* spheroplasts increased in volume when L-Ara was removed from the growth media, demonstrating that cell metabolism was very rapidly restored (Figure 3D). However, WigKR played an essential role in the recovery of L-Ara treated cells: the expansion in volume of the *wigKR* spherical cells upon L-Ara removal led to their lysis (Figure 3D). These results indicate that the recovery of a constitutive level of PG synthesis was not sufficient to expand the residual amount of cell wall left in the spheroplasts to accommodate the increase in cellular material, as it was observed for the spheroplasts induced by cell wall targeting antibiotics. Taken together, these results fit with the idea that the histidine kinase/response regulator pair *wigKR* serves to induce a higher expression of the full set of cell wall synthetic genes in response to cell wall damages [22].

### Spheroplast formation could be a general survival strategy of *Vibrios*

A multitude of *Vibrio* species, including *V. vulnificus*, *V. shiloi*, *V. tasmaniensis*, *V. parahaemolyticus* and *V. cholerae* form Viable But Not Culturable (VBNC) cells in cold temperatures [46–48]. Cold-induced VBNC cells are osmo-resistant and remain viable for months until the next warm season, which partly explains the seasonality of *Vibrio* epidemics [46–48]. Cold is supposed to perturb the metabolic activities of the above-mentioned species, plunging them in a state of so-called dormancy [49–51]. L-Ara induced spheroplasts are akin to cold-induced VBNC cells: their formation results from the perturbation of their metabolic activities by a by-product of L-Ara degradation (Figure 4); they are apparently dormant since they do not grow in volume (Figure 3); they are osmo-resistant and return to proliferation upon the removal of L-Ara (Figure 3). *Vibrios* have also been described to persist as coccoidal bodies in biofilms in the natural aquatic environment [4,52–54]. Interestingly, L-Ara was found to induce biofilm formation in *Vibrio fischeri*, even though no remarks were made about cell morphology [55]. As in *V. cholerae* spheroplasts, mutations in GalK or the galactose transporter suppressed the phenotypic action of L-Ara [55]. Taken together, these observations suggest that the effect of L-Ara on *V. cholerae* could serve as a paradigm for a general survival strategy of *Vibrio* species under a variety of environmental stresses.

## Methods

### Plasmids and strains

Bacterial strains and plasmids used in this study are listed in Supplementary Table 1. Strains were rendered competent by the insertion of *hapR* by specific transposition and constructed by natural transformation. Engineered strains were confirmed by PCR.

### Growth curves

Cells were grown at 30°C in M9 minimal medium supplemented with 0.2% (w/v) fructose and 1 μg/ml thiamine (M9-MM), M9-MM + 0.1% casamino acids (M9-MM + CAA) and Luria-Bertani broth (LB) in a 96-well microtitre plate and the optical density at 600 nm followed over time in a Tecan plate reader. The growth curves plotted are the average of three replicates; the standard deviation is represented for each time point. For CFU and rod to sphere kinetics, cells were grown in flasks in M9-MM at 30°C, 0.2% (w/v) L-Ara was added when indicated. Samples were taken every hour for plating and/or microscopic inspection. Three replicates were performed for each experiment.

### L-Ara survival assay

Over-night wild-type (EPV50) and *wigKR* deletion mutant (EGV515) cultures were diluted 200 times in M9-MM, followed by 2 hours of growth at 30°C before 0.2% (w/v) L-Ara was added. Cells were checked for transition to spherical morphology at the microscope. Serial dilutions of T_0_ (before L-Ara addition) and T_10_ (after L-Ara treatment) samples were plated on LB plates and the number of colonies used to calculate the CFU at T_0_ and T_10_. The ratio CFU T_10_ / CFU T_0_ is used to calculate the percentage of cells able to survive L-Ara treatment and revert to proliferation.

### Microscopy

Cells were spread on a 1% (w/v) agar pad (ultrapure agarose, Invitrogen) for analysis. For snapshots, images were acquired using a DM6000-B (Leica) microscope. For time-lapse analyses the agarose pad was made using M9-MM with 0.2% (w/v) L-Ara if needed and images were acquired using an Evolve 512 EMCCD camera (Roper Scientific) attached to an Axio Observe spinning disk (Zeiss). To observe rod to sphere transition on agarose pads, 0.2% (w/v) L-Ara was added in liquid M9-MM cultures 2 hours before transferring cells on agarose pads containing L-Ara and starting microscopic imaging.

### Transposon mutagenesis screen

The transposon mutagenesis was performed conjugating the *E. coli* strain SM10 λ *pir*/pSC189, which carries a mini-Himar transposon associated to a kanamycin resistance, with the *V. cholerae* strain EGV299. We constructed two Tn libraries of approximately 300,000 clones each. Tn mutants were selected on M9-MM plates containing kanamycin and 0.1% (w/v) L-Ara incubated overnight at 30°C. Mutants able to grow on plate were isolated and inspected at the microscope for growth and morphology in presence of L-Ara. An arbitrary PCR followed by DNA sequencing was performed to determine the transposon insertion site.

### Peptidoglycan analysis

EGV217 over-night cultures were diluted 200 times in M9-MM and grown at 30°C. 100 ml were centrifuged after 7 hours of growth and 1L of culture, to which 0.2% (w/v) L-Ara was added after 2 hours, was harvested after further 7 hours of incubation. Cells were checked for complete transition to spherical morphology at the microscope before harvesting. Previously described methods were followed for muropeptide isolation and ultra-performance liquid chromatography (UPLC) analysis [56,57]. After boiling for 2 hours cell pellets with SDS (sodium dodecyl sulfate) the lysates were left stirring over night at room temperature. Cell wall material was pelleted, washed with MQ water to remove the SDS, and digested with pronase E to remove Braun’s lipoprotein. Purified peptidoglycan was re-suspended in MQ water and treated over night with muramidase at 37°C. Soluble muropeptides were reduced with sodium borohydride and the pH then adjusted to 3.5 with phosphoric acid. Samples were injected in an UPLC system to obtain the muropeptide profiles. UPLC separation was performed on a Waters UPLC system equipped with an ACQUITY UPLC BEH C18 Column, 130 Å, 1.7 μm, 2.1 mm × 150 mm (Waters) and a dual wavelength absorbance detector using a linear gradient from buffer A (phosphate buffer 50 mM, pH 4.35) to buffer B (phosphate buffer 50 mM, pH 4.95, methanol 15% (v/v)) in a 28-min run with a 0.25 mL/min flow. Elution of muropeptides was detected at 204 nm. Identity of the peaks was assigned by comparison of the retention times and profiles to other chromatograms in which mass spectrometry data have been collected. The relative amounts of the muropeptides and the percentage of cross-linkage were calculated as described by Glauner *et al*. [19]. To estimate the amount of peptidoglycan per cell, the total area of the chromatogram was normalized to the OD of the culture. All values are the means of three independent experiments.

## Supplementary Figure legends

**Supplementary Figure 1.**
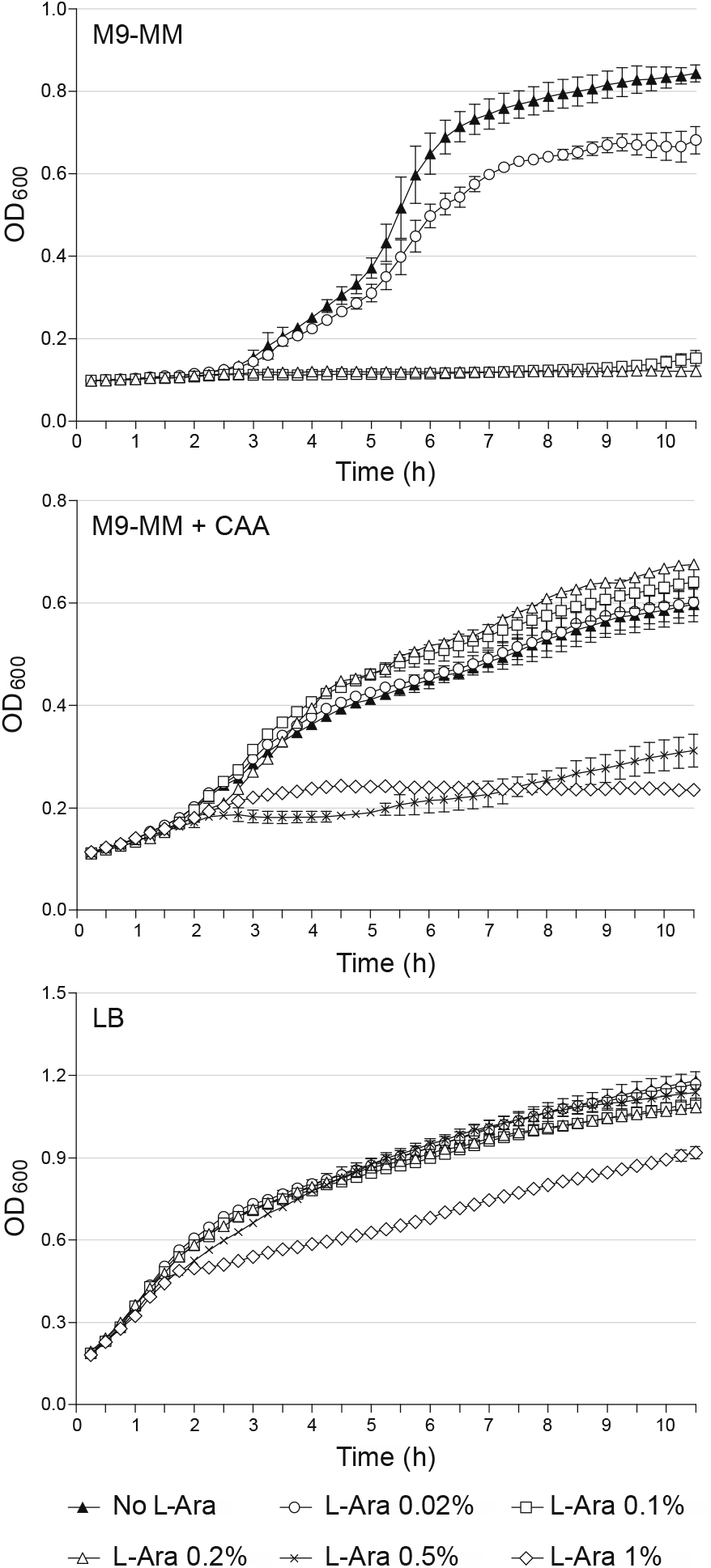
*V. cholerae* N16961 growth profiles in M9-MM (top panel), M9-MM + CAA (middle panel) and LB (bottom panel) in the presence of increasing concentrations of L-Ara. The optical density at 600 nm over time is the mean of three independent replicates; the standard deviation is represented for each time point.

**Supplementary Figure 2.**
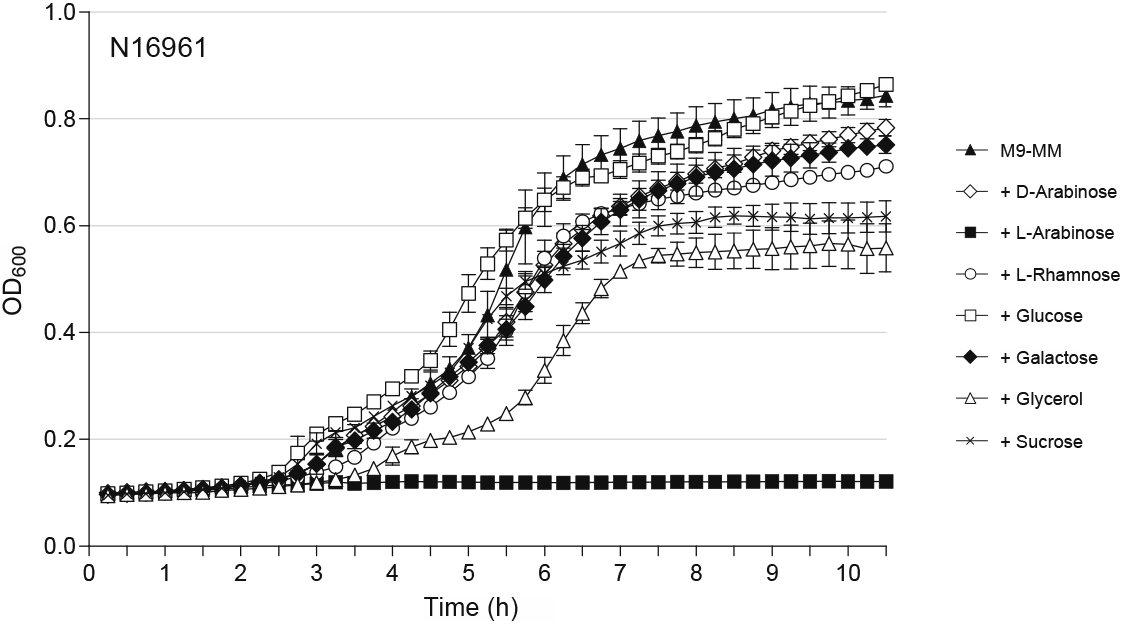
Carbon sources tested for inhibition of cell growth. *V. cholerae* N16961 growth profiles in M9-MM in the presence of different carbon sources at a concentration of 0.2% (w/v), with the exception of glycerol at 10 % (v/v). The optical density at 600 nm over time is the mean of three independent replicates; the standard deviation is represented for each time point.

**Supplementary Figure 3.**
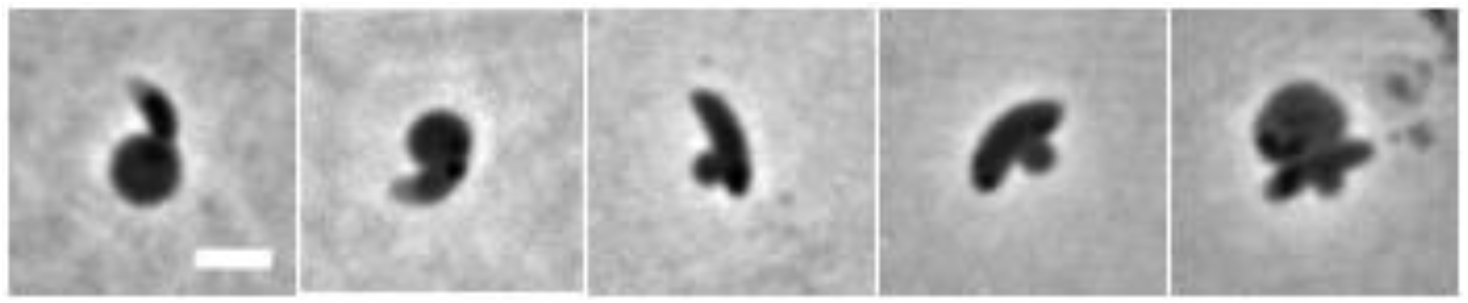
Localization of the bleb formation. N16961 cells were grown in M9-MM + 0.2% (w/v) L-Ara at 30°C. Phase contrast images. Scale bar = 2 μm.

**Supplementary Figure 4.**
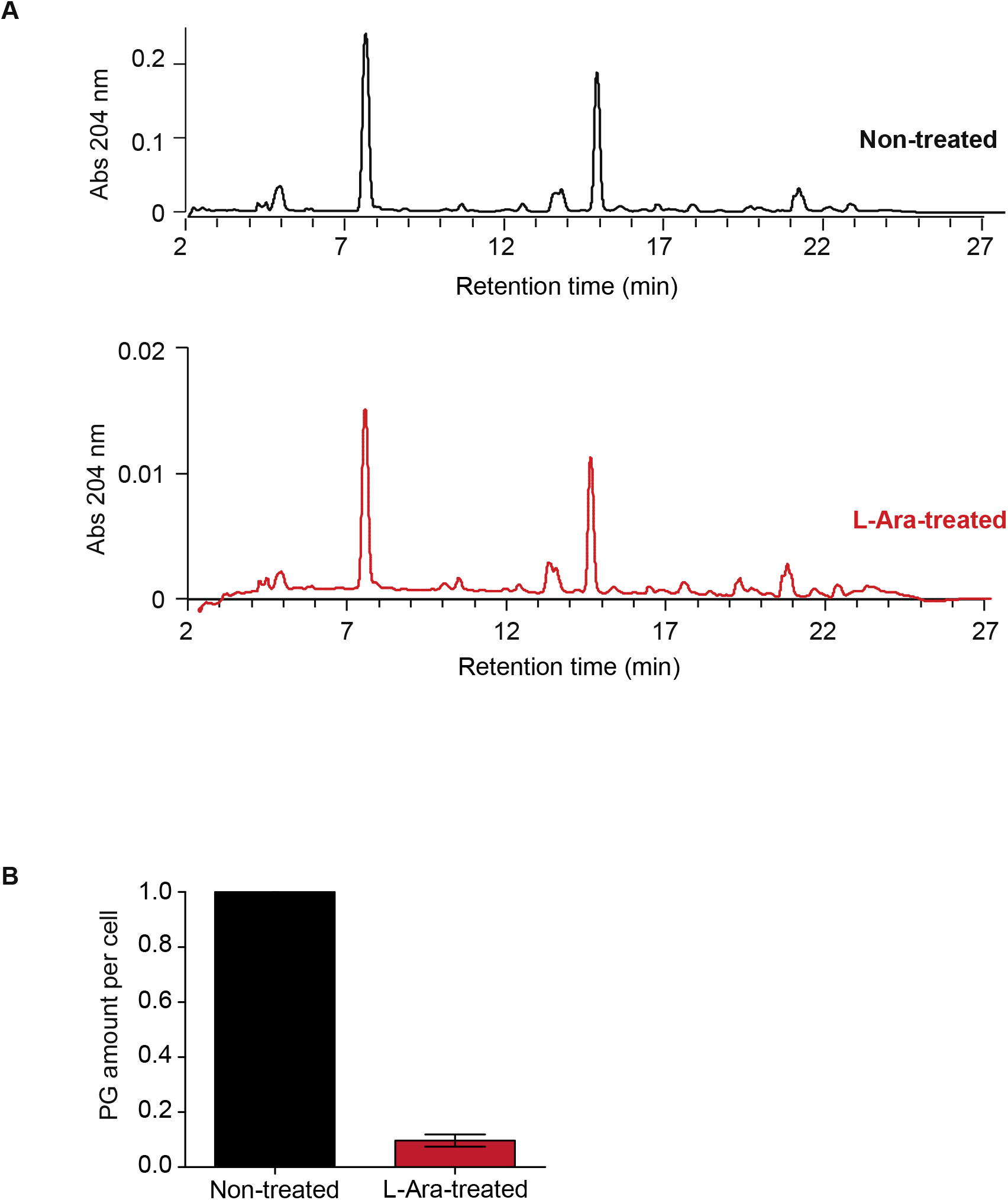
**A.** UPLC muropeptides separation. A representative muropeptide profile is presented for cells grown in M9-MM supplemented (red) or not (black) with 0.2% (w/v) L-Ara. **B.** Peptidoglycan quantification in non-treated and L-Ara treated cells. Amount of PG per cell normalized to the amount obtained for the non-treated cultures. All values are the means of three independent experiments. Mean of three independent replicates and the standard deviation are represented.

**Supplementary Figure 5.**
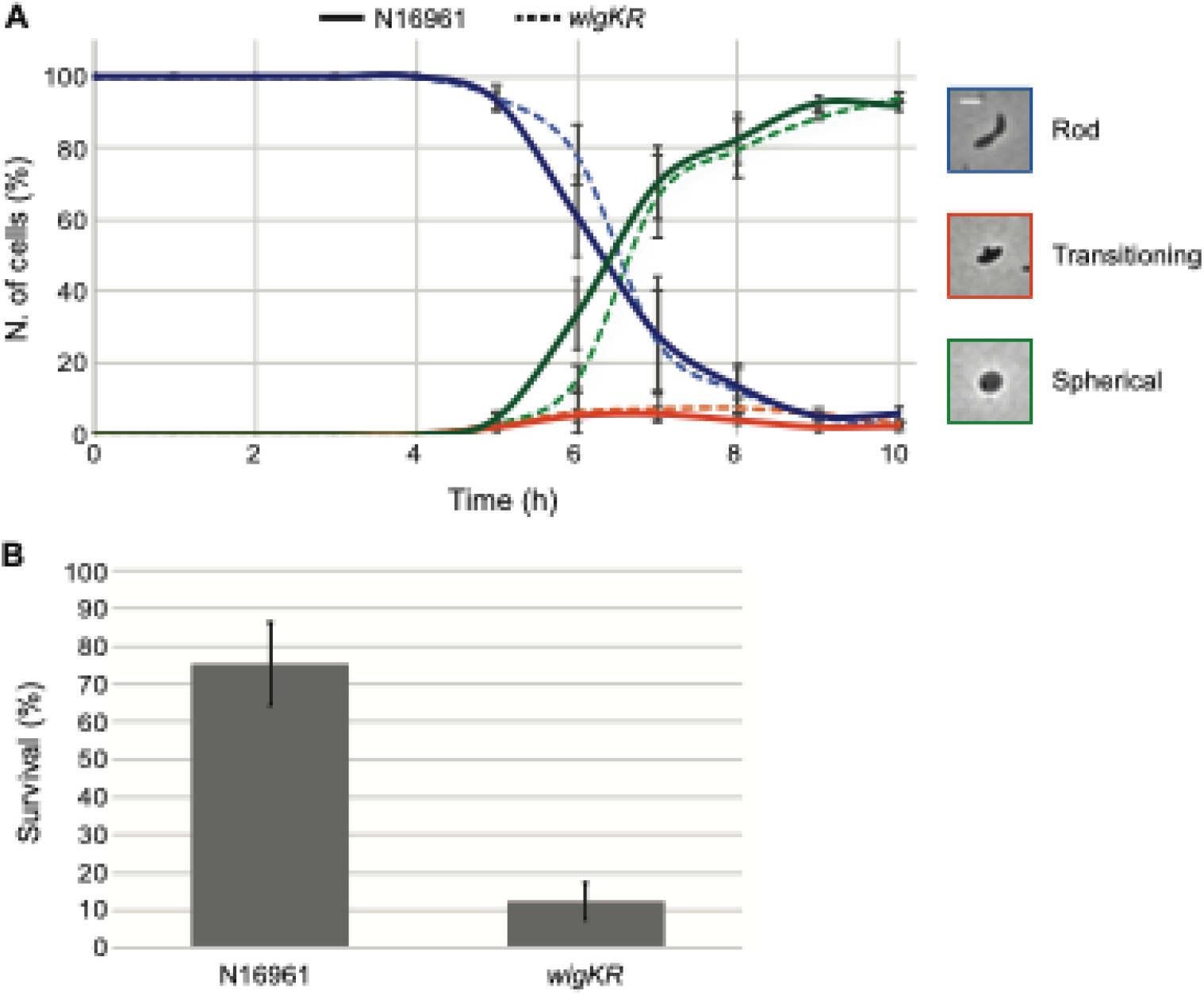
**A.** Morphological transition from rods to spheres of wild-type N16961 and *wigKR* mutant cells after addition of 0.2% (w/v) L-Ara. Cells were grown in M9-MM at 30°C and cell shape was inspected at the microscope every hour. A representative image for each cell category (rod, transitioning, spherical) is represented. Scale bar = 2 μm. **B.** Survival of wild-type N16961 and *wigKR* mutant cells after incubation in M9-MM + 0.2% (w/v) L-Ara for 10 hours. Data are averages of three biological replicates; the standard deviation is represented.

**Supplementary Figure 6.**
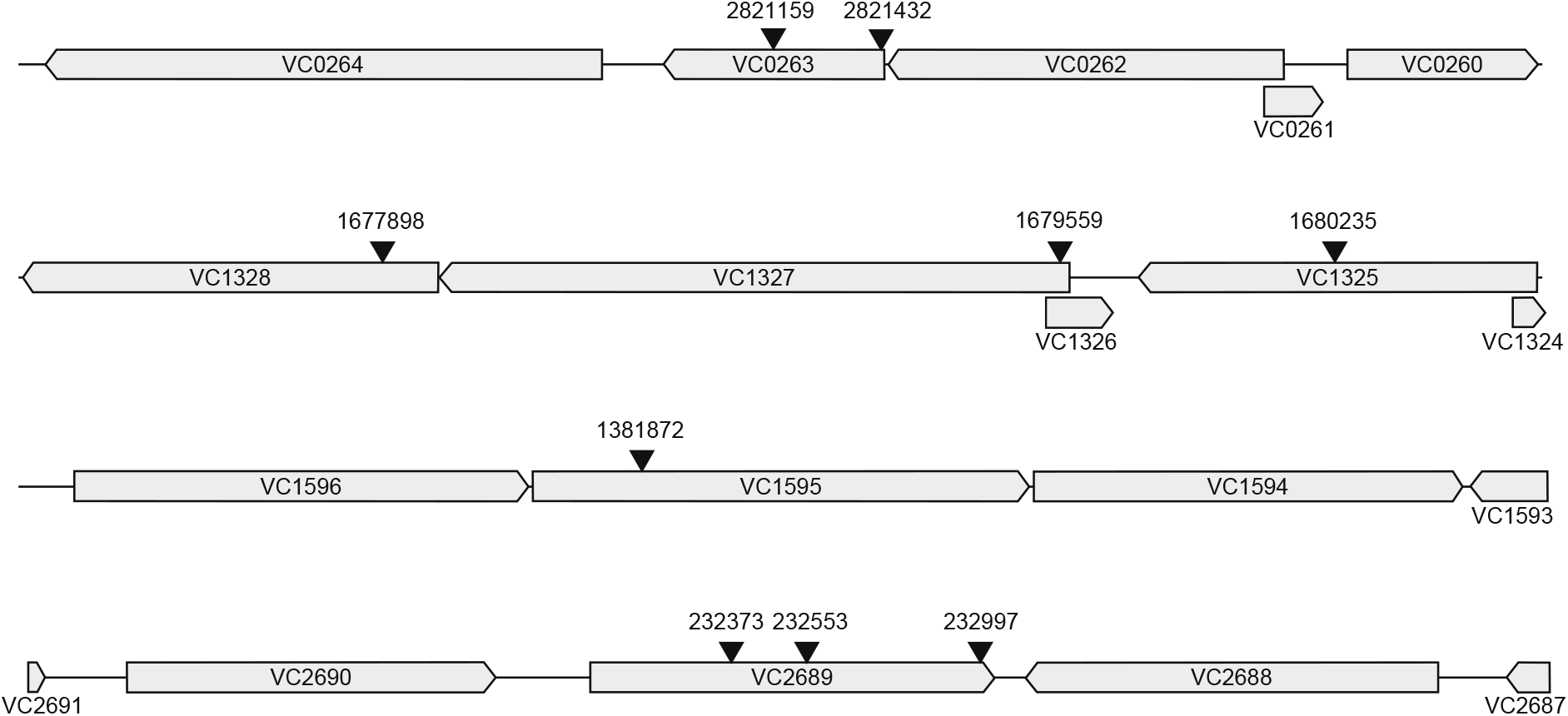
Maps of the regions of the chromosomes showing the location of Tn insertions in L-Ara insensitive mutants. The arrows represent the sites of insertion.

**Supplementary Figure 7.**
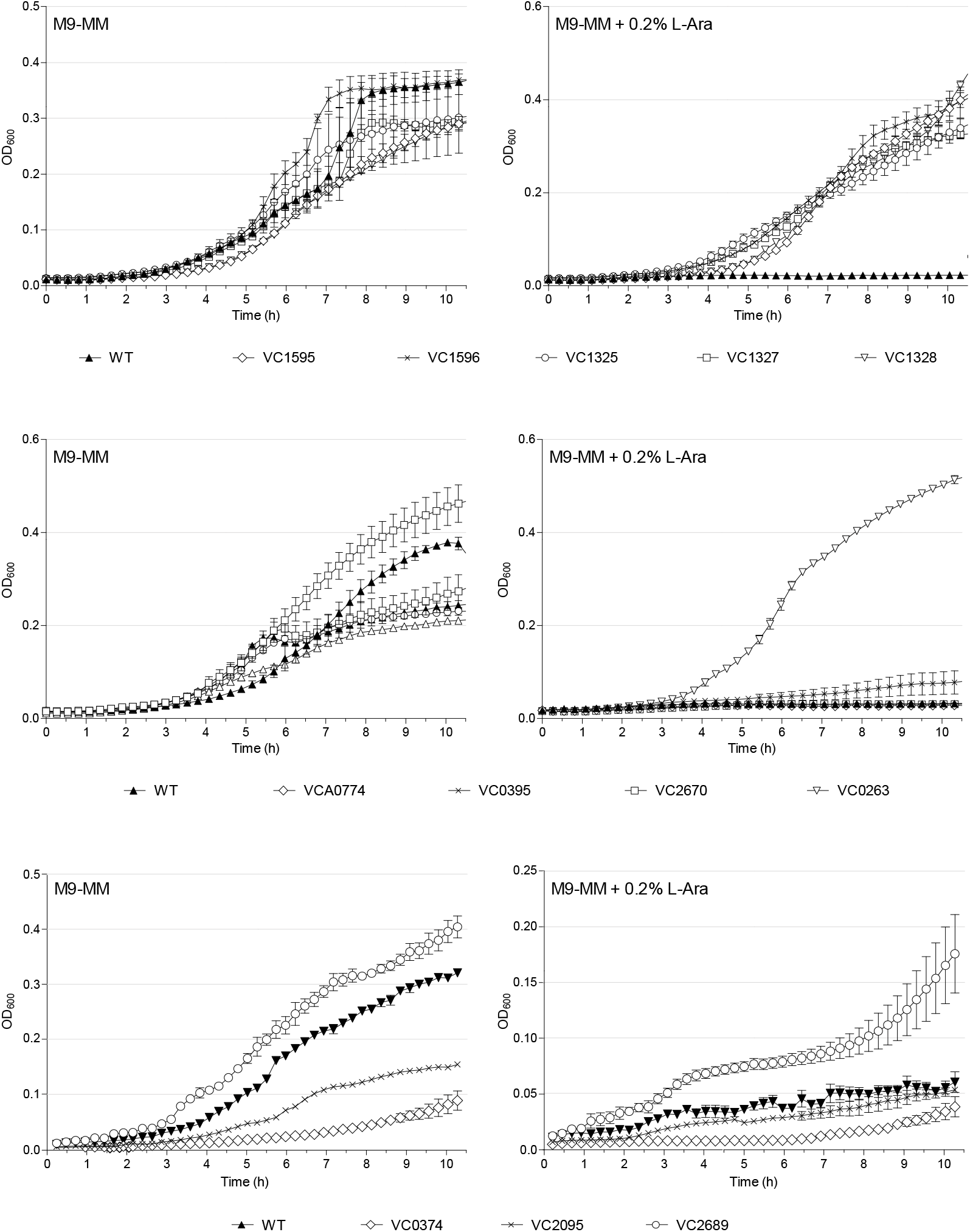
Growth profiles in M9-MM with and without 0.2% (w/v) L-Ara of *V. cholerae* strains carrying Tn inactivated mutants of genes belonging to the galactose Leloir and glycolysis pathways. The optical density at 600 nm over time is the mean of three independent replicates; the standard deviation is represented for each time point.

## Acknowledgements

We would like to acknowledge financial support from the Agence Nationale pour la Recherche [ANR19-CE35-0013-01 SurVi]. We thank C. Possoz for helpful discussions and Y. Yamaichi for providing the ordered *V. cholerae* mapped Tn library. Research in the Cava laboratory is supported by the Laboratory of Molecular Infection Medicine Sweden (MIMS), the Swedish Research Council (VR), the Knut and Alice Wallenberg Foundation (KAW) and the Kempe Foundation. S.B.H. was supported by a Martin Escudero Postdoctoral fellowship.

## References

1. The Biology of *Vibrios*. American Society of Microbiology; 2006. doi:10.1128/9781555815714

2. Chaiyanan S, Chaiyanan S, Grim C, Maugel T, Huq A, Colwell RR. Ultrastructure of coccoid viable but non-culturable *Vibrio cholerae*. Environmental Microbiology. 2007;9: 393–402. doi:10.1111/j.1462-2920.2006.01150.x

3. Huq A, Colwell RR, Rahman R, Ali A, Chowdhury MA, Parveen S, et al. Detection of *Vibrio cholerae* O1 in the aquatic environment by fluorescent-monoclonal antibody and culture methods. Appl Environ Microbiol. 1990;56: 2370–2373.

4. Alam M, Sultana M, Nair GB, Siddique AK, Hasan NA, Sack RB, et al. Viable but nonculturable *Vibrio cholerae* O1 in biofilms in the aquatic environment and their role in cholera transmission. Proc Natl Acad Sci USA. 2007;104: 17801–17806. doi:10.1073/pnas.0705599104

5. Dörr T, Davis BM, Waldor MK. Endopeptidase-mediated beta lactam tolerance. PLoS Pathog. 2015;11: e1004850. doi:10.1371/journal.ppat.1004850

6. Clemens JD, Nair GB, Ahmed T, Qadri F, Holmgren J. Cholera. Lancet. 2017;390: 1539–1549. doi:10.1016/S0140-6736(17)30559-7

7. Kim EJ, Lee D, Moon SH, Lee CH, Kim SJ, Lee JH, et al. Molecular Insights Into the Evolutionary Pathway of *Vibrio cholerae* O1 Atypical El Tor Variants. PLoS Pathog. 2014;10: e1004384. doi:10.1371/journal.ppat.1004384

8. Chun J, Grim CJ, Hasan NA, Lee JH, Choi SY, Haley BJ, et al. Comparative genomics reveals mechanism for short-term and long-term clonal transitions in pandemic *Vibrio cholerae*. Proc Natl Acad Sci U S A. 2009;106: 15442–7. doi:0907787106 [pii] 10.1073/pnas.0907787106

9. Weill F-X, Domman D, Njamkepo E, Tarr C, Rauzier J, Fawal N, et al. Genomic history of the seventh pandemic of cholera in Africa. Science. 2017;358: 785–789. doi:10.1126/science.aad5901

10. Domman D, Quilici M-L, Dorman MJ, Njamkepo E, Mutreja A, Mather AE, et al. Integrated view of *Vibrio cholerae* in the Americas. Science. 2017;358: 789–793. doi:10.1126/science.aao2136

11. Mutreja A, Kim DW, Thomson NR, Connor TR, Lee JH, Kariuki S, et al. Evidence for several waves of global transmission in the seventh cholera pandemic. Nature. 2011;477: 462–465. doi:10.1038/nature10392

12. Guzman LM, Belin D, Carson MJ, Beckwith J. Tight regulation, modulation, and high-level expression by vectors containing the arabinose P_BAD_ promoter. J Bacteriol. 1995;177: 4121–30.

13. Lee N, Francklyn C, Hamilton EP. Arabinose-induced binding of AraC protein to *araI2* activates the *araBAD* operon promoter. Proc Natl Acad Sci USA. 1987;84: 8814–8818.

14. Golder T, Mukhopadhyay AK, Koley H, Nandy RK. Nonmetabolizable arabinose inhibits *Vibrio cholerae* growth in M9 medium with gluconate as sole carbon source. Jpn J Infect Dis. 2020. doi:10.7883/yoken.JJID.2019.304

15. Oliver JD. Recent findings on the viable but nonculturable state in pathogenic bacteria. FEMS Microbiol Rev. 2010;34: 415–425. doi:10.1111/j.1574-6976.2009.00200.x

16. Bianco PR, Lyubchenko YL. SSB and the RecG DNA helicase: an intimate association to rescue a stalled replication fork. Protein Sci. 2017;26: 638–649. doi:10.1002/pro.3114

17. Galli E, Poidevin M, Le Bars R, Desfontaines J-M, Muresan L, Paly E, et al. Cell division licensing in the multi-chromosomal *Vibrio cholerae* bacterium. Nature Microbiology. 2016;1: 16094.

18. Allan EJ, Hoischen C, Gumpert J. Bacterial L-forms. Adv Appl Microbiol. 2009;68: 1–39. doi:10.1016/S0065-2164(09)01201-5

19. Glauner B, Höltje JV, Schwarz U. The composition of the murein of *Escherichia coli*. J Biol Chem. 1988;263: 10088–10095.

20. van Heijenoort J. Peptidoglycan hydrolases of *Escherichia coli*. Microbiol Mol Biol Rev. 2011;75: 636–663. doi:10.1128/MMBR.00022-11

21. Vollmer W, Joris B, Charlier P, Foster S. Bacterial peptidoglycan (murein) hydrolases. FEMS Microbiol Rev. 2008;32: 259–286. doi:10.1111/j.1574-6976.2007.00099.x

22. Dörr T, Alvarez L, Delgado F, Davis BM, Cava F, Waldor MK. A cell wall damage response mediated by a sensor kinase/response regulator pair enables beta-lactam tolerance. Proc Natl Acad Sci USA. 2016;113: 404–409. doi:10.1073/pnas.1520333113

23. Cameron DE, Urbach JM, Mekalanos JJ. A defined transposon mutant library and its use in identifying motility genes in *Vibrio cholerae*. Proc Natl Acad Sci USA. 2008;105: 8736–8741. doi:10.1073/pnas.0803281105

24. Harayama S, Bollinger J, Iino T, Hazelbauer GL. Characterization of the *mgl* operon of *Escherichia coli* by transposon mutagenesis and molecular cloning. J Bacteriol. 1983;153: 408–415.

25. Hogg RW, Voelker C, Von Carlowitz I. Nucleotide sequence and analysis of the *mgl* operon of *Escherichia coli* K12. Mol Gen Genet. 1991;229: 453–459.

26. Kalckar HM, Kurahashi K, Jordan E. Hereditary defects in galactose metabolism in *Escherichia coli* mutants I. determination of enzyme activities. Proc Natl Acad Sci USA. 1959;45: 1776–1786.

27. Stevenson G, Andrianopoulos K, Hobbs M, Reeves PR. Organization of the *Escherichia coli* K-12 gene cluster responsible for production of the extracellular polysaccharide colanic acid. J Bacteriol. 1996;178: 4885–4893. doi:10.1128/jb.178.16.4885-4893.1996

28. Patel KB, Toh E, Fernandez XB, Hanuszkiewicz A, Hardy GG, Brun YV, et al. Functional characterization of UDP-glucose:undecaprenyl-phosphate glucose-1-phosphate transferases of *Escherichia coli* and *Caulobacter crescentus*. J Bacteriol. 2012;194: 2646–2657. doi:10.1128/JB.06052-11

29. Geddes BA, Oresnik IJ. Inability to catabolize galactose leads to increased ability to compete for nodule occupancy in *Sinorhizobium meliloti*. J Bacteriol. 2012;194: 5044–5053. doi:10.1128/JB.00982-12

30. Frey PA. The Leloir pathway: a mechanistic imperative for three enzymes to change the stereochemical configuration of a single carbon in galactose. FASEB J. 1996;10: 461–470.

31. Shiba T, Hill RT, Straube WL, Colwell RR. Decrease in culturability of *Vibrio cholerae* caused by glucose. Appl Environ Microbiol. 1995;61: 2583–2588.

32. Englesberg E, Anderson RL, Weinberg R, Lee N, Hoffee P, Huttenhauer G, et al. L-Arabinose-sensitive, L-ribulose 5-phosphate 4-epimerase-deficient mutants of *Escherichia coli*. J Bacteriol. 1962;84: 137–146.

33. Kurahashi K, Wahba AJ. Interference with growth of certain *Escherichia coli* mutants by galactose. Biochim Biophys Acta. 1958;30: 298–302.

34. Yarmolinsky MB, Wiesmeyer H, Kalckar HM, Jordan E. Hereditary defects in galactose metabolism in *Escherichia coli* mutants II. Galactose-induced sensitivity. Proc Natl Acad Sci USA. 1959;45: 1786–1791.

35. Englesberg E, Baron LS. Mutation to L-rhamnose resistance and transduction to L-rhamnose utilization in *Salmonella typhosa*. J Bacteriol. 1959;78: 675–686.

36. Weaver AI, Murphy SG, Umans BD, Tallavajhala S, Onyekwere I, Wittels S, et al. Genetic Determinants of Penicillin Tolerance in *Vibrio cholerae*. Antimicrob Agents Chemother. 2018;62. doi:10.1128/AAC.01326-18

37. Cross T, Ransegnola B, Shin J-H, Weaver A, Fauntleroy K, VanNieuwenhze MS, et al. Spheroplast-Mediated Carbapenem Tolerance in Gram-Negative Pathogens. Antimicrob Agents Chemother. 2019;63. doi:10.1128/AAC.00756-19

38. Ranjit DK, Young KD. The Rcs stress response and accessory envelope proteins are required for de novo generation of cell shape in *Escherichia coli*. J Bacteriol. 2013;195: 2452–2462. doi:10.1128/JB.00160-13

39. Joseleau-Petit D, Liébart J-C, Ayala JA, D’Ari R. Unstable *Escherichia coli* L forms revisited: growth requires peptidoglycan synthesis. J Bacteriol. 2007;189: 6512–6520. doi:10.1128/JB.00273-07

40. Cambré A, Zimmermann M, Sauer U, Vivijs B, Cenens W, Michiels CW, et al. Metabolite profiling and peptidoglycan analysis of transient cell wall-deficient bacteria in a new *Escherichia coli* model system. Environ Microbiol. 2015;17: 1586–1599. doi:10.1111/1462-2920.12594

41. Mercier R, Kawai Y, Errington J. General principles for the formation and proliferation of a wall-free (L-form) state in bacteria. Elife. 2014;3. doi:10.7554/eLife.04629

42. Magnet S, Dubost L, Marie A, Arthur M, Gutmann L. Identification of the L,D-transpeptidases for peptidoglycan cross-linking in *Escherichia coli*. J Bacteriol. 2008;190: 4782–4785. doi:10.1128/JB.00025-08

43. Cava F, de Pedro MA, Lam H, Davis BM, Waldor MK. Distinct pathways for modification of the bacterial cell wall by non-canonical D-amino acids. EMBO J. 2011;30: 3442–3453. doi:10.1038/emboj.2011.246

44. Peters NT, Dinh T, Bernhardt TG. A fail-safe mechanism in the septal ring assembly pathway generated by the sequential recruitment of cell separation amidases and their activators. J Bacteriol. 2011;193: 4973–4983. doi:10.1128/JB.00316-11

45. Möll A, Dörr T, Alvarez L, Chao MC, Davis BM, Cava F, et al. Cell separation in *Vibrio cholerae* is mediated by a single amidase whose action is modulated by two nonredundant activators. J Bacteriol. 2014;196: 3937–3948. doi:10.1128/JB.02094-14

46. Alam M, Sultana M, Nair GB, Sack RB, Sack DA, Siddique AK, et al. Toxigenic *Vibrio cholerae* in the aquatic environment of Mathbaria, Bangladesh. Appl Environ Microbiol. 2006;72: 2849–2855. doi:10.1128/AEM.72.4.2849-2855.2006

47. Huq A, Colwell RR, Rahman R, Ali A, Chowdhury MA, Parveen S, et al. Detection of *Vibrio cholerae* O1 in the aquatic environment by fluorescent-monoclonal antibody and culture methods. Appl Environ Microbiol. 1990;56: 2370–2373. Available: https://aem.asm.org/content/56/8/2370

48. Su C-P, Jane W-N, Wong H. Changes of ultrastructure and stress tolerance of *Vibrio parahaemolyticus* upon entering viable but nonculturable state. Int J Food Microbiol. 2013;160: 360–366. doi:10.1016/j.ijfoodmicro.2012.11.012

49. Rittershaus ESC, Baek S-H, Sassetti CM. The normalcy of dormancy: common themes in microbial quiescence. Cell Host Microbe. 2013;13: 643–651. doi:10.1016/j.chom.2013.05.012

50. Lewis K. Persister cells. Annu Rev Microbiol. 2010;64: 357–372. doi:10.1146/annurev.micro.112408.134306

51. Hayes CS, Low DA. Signals of growth regulation in bacteria. Curr Opin Microbiol. 2009;12: 667–673. doi:10.1016/j.mib.2009.09.006

52. Rodrigues S, Paillard C, Le Pennec G, Dufour A, Bazire A. *Vibrio tapetis*, the Causative Agent of Brown Ring Disease, Forms Biofilms with Spherical Components. Front Microbiol. 2015;6: 1384. doi:10.3389/fmicb.2015.01384

53. Oliver JD, Hite F, McDougald D, Andon NL, Simpson LM. Entry into, and resuscitation from, the viable but nonculturable state by *Vibrio vulnificus* in an estuarine environment. Appl Environ Microbiol. 1995;61: 2624–2630.

54. Grimes DJ, Atwell RW, Brayton PR, Palmer LM, Rollins DM, Roszak DB, et al. The fate of enteric pathogenic bacteria in estuarine and marine environments. Microbiol Sci. 1986;3: 324–329.

55. Visick KL, Quirke KP, McEwen SM. Arabinose induces pellicle formation by *Vibrio fischeri*. Appl Environ Microbiol. 2013;79: 2069–2080. doi:10.1128/AEM.03526-12

56. Desmarais SM, De Pedro MA, Cava F, Huang KC. Peptidoglycan at its peaks: how chromatographic analyses can reveal bacterial cell wall structure and assembly. Mol Microbiol. 2013;89: 1–13. doi:10.1111/mmi.12266

57. Möll A, Dörr T, Alvarez L, Davis BM, Cava F, Waldor MK. A D, D-carboxypeptidase is required for *Vibrio cholerae* halotolerance. Environ Microbiol. 2015;17: 527–540. doi:10.1111/1462-2920.12779

